# Integrated transcriptomics and metabolomics reveal signatures of lipid metabolism dysregulation in HepaRG liver cells exposed to PCB 126

**DOI:** 10.1101/259093

**Authors:** Robin Mesnage, Martina Biserni, Sucharitha Balu, Clément Frainay, Nathalie Poupin, Fabien Jourdan, Eva Wozniak, Theodoros Xenakis, Charles A Mein, Michael N Antoniou

## Abstract

Chemical pollutant exposure is a risk factor contributing to the growing epidemic of nonalcoholic fatty liver disease (NAFLD) affecting human populations that consume a Western diet. Although it is recognized that intoxication by chemical pollutants can lead to NAFLD, there is limited information available regarding the mechanism by which typical environmental levels of exposure can contribute to the onset of this disease. Here we describe the alterations in gene expression profiles and metabolite levels in the human hepatocyte HepaRG cell line, a validated model for cellular steatosis, exposed to the polychlorinated biphenyl (PCB) 126, one of the most potent chemical pollutants that can induce NAFLD. Sparse partial least squares classification of the molecular profiles revealed that exposure to PCB 126 provoked a decrease in polyunsaturated fatty acids as well as an increase in sphingolipid levels, concomitant with a decrease in the activity of genes involved in lipid metabolism. This was associated with an increased oxidative stress reflected by marked disturbances in taurine metabolism. A gene ontology analysis showed hallmarks of an activation of the AhR receptor by dioxin-like compounds. These changes in metabolome and transcriptome profiles were observed even at the lowest concentration (100 pM) of PCB 126 tested. A decrease in docosatrienoate levels was the most sensitive biomarker. Overall, our integrated multi-omics analysis provides mechanistic insight into how this class of chemical pollutant can cause NAFLD. Our study lays the foundation for the development of molecular signatures of toxic effects of chemicals causing fatty liver diseases to move away from a chemical risk assessment based on *in vivo* animal experiments.

## Introduction

A growing epidemic of non-alcoholic fatty liver disease (NAFLD) is affecting human populations that consume a Western diet (Argyrou et al., 2017). The spectrum of NAFLD ranges from fatty infiltration of the liver (steatosis), to more advanced stages characterized by liver inflammation and fibrosis (non-alcoholic steatohepatitis, NASH), and subsequent cirrhosis, and hepatocellular carcinoma (Michelotti et al., 2013; Vernon et al., 2011). NAFLD currently affects 25% of the US population, and approximately 2–5% have NASH (Vernon et al., 2011) with concomitant massive clinical and economic burden (Younossi et al., 2016). Annual direct medical costs have been estimated to be approximately $103 billion ($1,613 per patient) in the US. The occurrence of NAFLD in Europe is also very high and ranges from 20-30% with annual medical costs currently running at €35 billion (from €354 to €1,163 per patient) in countries such as Germany, France, Italy, and the United Kingdom (Younossi et al., 2016). Risk factors identified to date for the development of NAFLD include being overweight or obese, having diabetes and high cholesterol or high triglycerides in the blood. Rapid weight loss and poor eating habits can also lead to NAFLD. The heritability of NAFLD was estimated at ~50% (Sookoian and Pirola, 2017). However, some individuals develop NAFLD even if they do not have any of these risk factors (Foulds et al., 2017) and with exposure to physiologically active environmental pollutants being a recognised additional determinant of disease establishment.

Chemical exposure has long been known to be a risk factor for NAFLD/NASH (Wahlang et al., 2013), which is also known as toxicant-associated fatty liver disease (TAFLD) and its more, severe form, toxicant-associated steatohepatitis (TASH). TAFLD has been commonly associated with acute chemical exposures at the workplace (Cave et al., 2010b), or after industrial accidents (Yu et al., 1997). However, there is also evidence showing that environmental pollutants have the ability to cause TAFLD at environmental levels of exposure (Cave et al., 2010a). The ability of environmental pollutants to disturb metabolic processes at low environmental levels of exposure generally results from their ability to interfere with the activity of natural hormones by binding to their receptors (Heindel et al., 2017). Different structural classes of environmental pollutants have been shown to modulate liver nuclear receptors, such as dioxins acting on the aryl hydrocarbon receptor (AhR), organotins on peroxisome proliferator-activated receptor γ (PPARγ), or organochlorines on the constitutive androstane receptor (CAR) (Foulds et al., 2017). Some environmental pollutants also promote NAFLD because of their hormone mimicking properties on distant organs. For example, bisphenol A acts on the pancreas by stimulating the production and secretion of insulin which, in turn, increases the storage of fat by stimulating lipogenesis in the liver (Fabricio et al., 2016). Commonly defined as obesogens, some other endocrine disrupting chemicals (EDCs) have been found to promote adipose cell differentiation and lipid accumulation (Janesick and Blumberg, 2016) with some studies establishing a direct link between human exposures to some EDCs and risk of obesity (Tang-Peronard et al., 2014). Recent advances in human microbial ecology has also revealed that alterations of the gut microbiota by chemicals at early stages of life accelerates hepatic lipid metabolism (Ba et al., 2017). However, TAFLD is not only a hormone receptor-mediated disease but can also arise from metabolic imbalances. For instance, fatty acid breakdown can be altered following a direct inhibition of mitochondrial β-oxidation by, for example, the pesticide chlordecone (Kaiser et al., 2012). Mitochondrial β-oxidation can also be indirectly inhibited by an alteration in NAD+ cofactor metabolism by compounds such as vinyl chloride.

Overall, biological mechanisms underlying the development of steatosis are grouped into 4 biological processes affecting the metabolism of fatty acids: an increased uptake, a decreased efflux, an increased synthesis, or defects in their oxidative metabolism (Angrish et al., 2017). All these processes are interrelated with a need for more studies providing mechanistic insight into the alteration of hepatic lipid metabolism, which in turn induces the development of NAFLD (Foulds et al., 2017). This is a priority for regulatory toxicology programs and includes a move away from a chemical risk assessment based on *in vivo* animal experiments (Merrick et al., 2015). This can be achieved by detailed molecular profiling studies describing the effects of known toxicants on validated tissue culture model systems, such as the human hepatocyte HepaRG cell line (Angrish et al., 2017).

We describe here the alterations in the gene expression profile and metabolite levels of HepaRG cells exposed to the polychlorinated biphenyl (PCB) congener 126, one of the most potent chemicals associated with development of TAFLD (Al-Eryani et al., 2015). Our investigation included a metabolomics analysis, which identified 802 metabolites, which were analysed with MetExplore (Cottret et al., 2010), in order to extract the metabolic sub-network involved in the biological response to low environmental levels of PCB 126. This was coupled with a transcriptomics analysis allowing identification of gene networks involved in the response to PCB 126. Our study is the first in depth investigation of the molecular profiles underlying the toxic effects of PCB 126, integrating the transcriptome and the metabolome in HepaRG cells in response and ultimately highlighted TAFLD-associated sensitive biomarkers of exposure to this class of pollutant.

## Material and methods

### Reagents

All reagents and chemicals, unless otherwise specified, were of analytical grade and were purchased from Sigma-Aldrich (Gillingham, Dorset, UK). The PCB 126 (98.5% purity, CAS Number 57465-28-8) was purchased from LGC Standards GmbH (Wesel, Germany). Stock solutions of PCB 126 were made by dissolving in DMSO. The Williams’E medium + GlutaMAX^™^ were purchased from Gibco (Thermo Fisher, Loughborough, UK). The supplement ADD670, as well as the DMSO, was provided by Biopredic International (Rennes, France).

### HepaRG cell culture

HepaRG^™^ cells (HPR 116) were purchased from Biopredic International (Rennes, France). Cells were thawed, suspended and plated in the general purpose medium (Williams’E medium + GlutaMAX^™^) containing the ADD670 supplement. A total of 72,000 and 2,000,000 cells were plated in collagen-coated 96 well-plates (Greiner Bio-One, Germany) and 6 well-plates (Biopredic) respectively. All cells were cultured at 37°C and a 5% CO_2_ atmosphere. The medium was refreshed at days 2, 5 and 7 after initial plating. The cells were kept in the general purpose medium until day 8, when the culture becomes well organized and includes well-delineated trabeculae and many canaliculi-like structures. At this time, the culture is composed of primitive biliary epithelial cells and mature hepatocytes with basal metabolic activities similar to fresh hepatocyte cells. From day 8 to day 14, cells were switched to the test medium composed of Williams’E medium + GlutaMAX^™^ supplemented with 2% fetal bovine serum (FBS; GE Healthcare Life Sciences, Buckinghamshire, UK), 2mM L-glutamine (GE Healthcare Life Sciences), 10 µg/ml penicillin/streptomycin (Life technologies) and 1% DMSO, as well as different concentrations of PCB 126 or the solvent control. Three concentrations of the PCB were tested in order to cover a wide range of biological effects, starting from low environmental exposures (100 pM) to high concentrations of (1 uM), with an intermediate concentration (10 nM).

### Mass spectrometry-based metabolomics

Approximately 5,000,000 cells per sample were harvested from the 6 well-plate cultures to obtain a sufficient amount of material to perform the metabolomics analysis. Cells were detached using 0.05% trypsin EDTA (Fisher Scientific, Loughborough, UK), and collected by centrifugation in order to eliminate trypsin residues and cell pellets frozen, The frozen cell pellets were then sent to Metabolon Inc. (Durham, NC, USA) who conducted the metabolomics analysis. Samples were prepared using the automated MicroLab STAR^®^ system from Hamilton Company. Proteins were precipitated with methanol under vigorous shaking for 2 min (Glen Mills GenoGrinder 2000), followed by centrifugation. Samples were placed briefly on a TurboVap^®^ (Zymark) to remove the organic solvent. The sample extracts were stored overnight under nitrogen before preparation for analysis. The resulting extract was analysed on four independent instrument platforms: two different separate reverse phase ultrahigh performance liquid chromatography-tandem mass spectroscopy analysis (RP/UPLC-MS/MS) with positive ion mode electrospray ionization (ESI), a RP/UPLC-MS/MS with negative ion mode ESI, as well as a by hydrophilic-interaction chromatography (HILIC)/UPLC-MS/MS with negative ion mode ESI.

All UPLC-MS/MS methods utilized a Waters ACQUITY ultra-performance liquid chromatography (UPLC) and a Thermo Scientific Q-Exactive high resolution/accurate mass spectrometer interfaced with a heated electrospray ionization (HESI-II) source and Orbitrap mass analyzer operated at 35,000 mass resolution. The sample extract was dried then reconstituted in solvents compatible to each of four methods used. Each reconstitution solvent contained a series of standards at fixed concentrations to ensure injection and chromatographic consistency. One aliquot was analyzed using acidic positive ion conditions, chromatographically optimized for more hydrophilic compounds. In this method, the extract was gradient eluted from a C18 column (Waters UPLC BEH C18-2.1x100 mm, 1.7 µm) using water and methanol, containing 0.05% perfluoropentanoic acid (PFPA) and 0.1% formic acid (FA). Another aliquot was also analyzed using acidic positive ion conditions, chromatographically optimized for more hydrophobic compounds. In this method, the extract was gradient eluted from the same afore mentioned C18 column using methanol, acetonitrile, water, 0.05% PFPA and 0.01% FA and was operated at an overall higher organic content. Another aliquot was analyzed using basic negative ion optimized conditions using a separate dedicated C18 column. The basic extracts were gradient eluted from the column using methanol and water, with 6.5mM ammonium bicarbonate at pH 8. The fourth aliquot was analyzed via negative ionization following elution from a HILIC column (Waters UPLC BEH Amide 2.1x150 mm, 1.7 µm) using a gradient consisting of water and acetonitrile with 10mM ammonium formate, pH 10.8. The MS analysis alternated between MS and data-dependent MSn scans using dynamic exclusion. The scan range varied slightly between methods but covered 70-1000 m/z.

Instrument variability was 5%. This was determined by calculating the median relative standard deviation (RSD) for the standards that were added to each sample prior to injection into the mass spectrometers. Overall process variability was 10%. This was determined by calculating the median RSD for all endogenous metabolites (that is, non-instrument standards) present in 100% of the pooled matrix samples.

Raw data was extracted, peak-identified and QC processed using Metabolon’s hardware and software. Compounds were identified by comparison to library entries of purified standards or recurrent unknown entities. Biochemical identifications are based on three criteria: retention index within a narrow retention time/index (RI) window of the proposed identification, accurate mass match to the library +/- 10 ppm, and the MS/MS forward and reverse scores between the experimental data and authentic standards. While there may be similarities between these molecules based on one of these factors, the use of all three data points can be utilized to distinguish and differentiate biochemicals. Peaks were quantified using area-under-the-curve.

### RNA Sequencing (RNA-seq)

#### RNA extraction and double-stranded cDNA library preparation

RNA extraction was performed using the Qiagen RNeasy kit according to the manufacturer’s instructions. The samples were checked for RNA quality using the Agilent 2100 Bioanalyzer and quantified using a Nanodrop instrument (ND-1000 Spectrophotometer; Thermo Fisher Scientific, Wilmington, DE, USA). The extracted RNA was subsequently subjected to treatment with DNAse I (Thermo Fisher Scientific Cat. No: AM222) followed by purification with Agencourt Ampure XP beads (Thermo Fisher Scientific Cat. No: A63881). A 20 ng aliquot of intact high quality total RNA (RIN>9) from each sample was then used as input to generate libraries for RNA-seq using the NEBNext Ultra II Directional RNA kit (NEB, Cat. no: E7760S) following the manufacturer’s recommendations. This protocol involved an initial step of rRNA depletion, followed by fragmentation prior to first strand cDNA synthesis and barcoding of the second-strand cDNA synthesised with indices for Illumina platform sequencing for final library amplification. The resulting libraries were assessed on the Bioanalyzer 2100 for purity. Sequencing of the resultant doublestranded (ds) cDNA libraries was performed by applying Illumina sequencing by synthesis technology.

#### Sequencing of ds-cDNA libraries

Out of a total of 40 cDNA libraries generated, 15 were randomly selected and assessed for quality and fragment size distribution using the Agilent 2200 Tapestation (Agilent Technologies, Waldbronn, Germany) prior to sequencing. All samples showed libraries of good quality, with minimal adapter-dimer contamination, and an average fragment size of 345bp. All 40 libraries were quantified using the Qubit 2.0 spectrophotometer and the high-sensitivity dsDNA Qubit reagent kit (Life Technologies, California, USA). An equimolar quantity of each library was pooled for sequencing, and the resulting complete pool was quality checked using the Agilent 2200 Tapestation. Sequencing consisted of generating 75bp paired-end reads for the final pool of 40 libraries using the Illumina NextSeq^®^500 instrument in conjunction with the NextSeq^®^500 v2 High-ouput 150- cycle kit (Illumina Inc., Cambridge, UK).

### Statistical analysis

The metabolome data analysis was performed using MetaboAnalyst 3.0 (Xia and Wishart, 2016). We removed variables with more than 50% of missing values. We also replaced remaining missing values with a small value (half of the minimum positive values in the original data) assuming to be the detection limit. Furthermore, variables that show low repeatability as measured using the relative standard deviation were removed. This step is recommended for untargeted metabolomics datasets with a large number of variables to remove baseline noise (Hackstadt and Hess, 2009). Data was then median scaled, log transformed, and normalized to protein concentration, as determined using the Bradford reagent, from the cell pellets. A principal component analysis (PCA) was first performed in order to inspect the data variance and see how the samples are related to each other. This was then followed by an orthogonal projection to latent structures discriminant analysis (OPLS-DA) to identify the source of variation between control and treated groups (Worley and Powers, 2016). OPLS-DA is an extension of the PLS-DA method, which incorporates an orthogonal component distinguishing the variability corresponding to the experimental perturbation (here the effect of PCB 126) from the portion of the data which is orthogonal; that is, independent from the experimental perturbation. Although PCA is an unsupervised method, the OPLS-DA is supervised. The separation between the different classes is calculated by maximizing the co-variance between the metabolome matrix (x) and the experimental group labels (y). It is thus prone to overfitting, which we addressed by performing permutation testing. Statistically significant alterations in the levels of metabolites were identified based on the significance after analysis with a one-way ANOVA test adjusted for multiple comparison with Fisher’s least significant difference (q<0.05). The metabolome network visualisation was performed with MetExplore as described (Cottret et al., 2010). Briefly, the sub-network was obtained by keeping reactions belonging to shortest paths up to a length of 4 between pairs of metabolites in the fingerprint. Network was modelled based on Recon2 (Thiele et al., 2013) human metabolic network by using atom conservation graph (Frainay and Jourdan, 2017) where two metabolites are connected if they are substrate and product of the same reaction and share at least one carbon (excluding carbon dioxide from the network).

The RNA-seq data analysis was performed using the latest version of the Tuxedo protocol with HISAT, StringTie and Ballgown (Pertea et al., 2016). First, we analysed the quality scores and other metrics using FASTQC (Andrews, 2017). Contamination from rRNA was measured by aligning the sequences of human rRNA to the FASTQ files (http://genomespot.blogspot.co.uk/2015/08/screen-for-rrna-contamination-in-rna.html). Sequences were then aligned to the human genome (Supplementary File 1) using the hierarchical indexing for spliced alignment of transcripts program HISAT2 (Kim et al., 2015). For this purpose, we used prebuilt indexes (H. sapiens, GRCh38) downloaded from the HISAT2 website. Alignment rates ranged from 85.12% to 95.53% (average of 91.63%). The number of reads per samples ranged from 9,467,558 to 35,623,876 (average of 21,575,770 reads). Then, StringTie was used to assemble and quantify the transcripts in each sample using the Homo_sapiens.GRCh38.89 database (Pertea et al., 2016). Finally, the differential gene expression analysis was made with Ballgown (Frazee et al., 2015) in R environment (Team., 2017). Low abundance transcripts with a variance across samples of less than one were filtered. A standard linear model-based comparison of transcript abundance was performed without adjusting for other covariates to identify differentially expressed transcripts (q<0.05).

Functional implications of the alteration in gene expression profiles were analysed using ClueGO (Bindea et al., 2009) and CluePedia plugins in Cytoscape (version 3.5.1)(Shannon et al., 2003). The GO biological process database (23.02.2017) and the KEGG annotation database (01.03.2017) were used. The analysis was conducted using a two-sided hypergeometric test for enrichment using a p-value threshold of 0.05 after its adjustment by the Benjamini-Hochberg procedure (Benjamini and Hochberg, 1995). GO term fusion was employed to integrate GO categories, minimize the output, and create a functionally organized GO category network. Our network displayed GO terms found in the levels 5-10 of the GO hierarchy, in order to avoid meaningless high-level hierarchy terms. Since it is well known that gene-annotation enrichment tools can give different results, we corroborated our interpretations by undertaking an additional KEGG functional analysis using the Wallenius non-central hypergeometric distribution in the Bioconductor package GOSeq (Young et al., 2010). This analysis has the advantage of correcting for the gene selection bias due to differences in transcript length present in RNA-seq datasets. KEGG annotation enrichment profiles were found to be very similar to the results obtained with ClueGO. These RNA-seq data have been submitted to Gene Omnibus and are accessible through accession number GSE109565.

The metabolome and transcriptome datasets were then integrated by a sparse Partial Least Squares regression (sPLS) performed with the MixOmics package. PLS is an supervised method, which selects correlated variables (genes, metabolites) across the same samples by maximizing the covariance between the datasets. The sparse version of PLS (sPLS) was used to select the most correlated variables using LASSO penalization on the pair of loading vectors as described (Le Cao et al., 2008).

## Results

We determined transcriptome and metabolome signatures of the effect of a 10-day exposure to PCB 126 in differentiated HepaRG cells (Figure 1). The cells presented no visible signs of aging at this time point. The metabolome platform detected 802 metabolites. We removed 7 variables with more than 50% missing values, and 199 variables that showed low repeatability. The effects of the PCB 126 were first visualized by plotting each sample as a point in space defined by the two principal components from a PCA, in order to reduce the 596-dimensional space defined by the variations in the metabolites levels (Figure 2A). The components separated the group of samples treated with the PCB126 from the control group. As the dose of PCB 126 increases, the groups become more clustered. We then built an OPLS-DA model (Figure 2B) on the basis of the PCA results. The OPLS-DA model for PCB 126 (R2X = 0.177, R2Y = 0.769, and Q2 = 0.58) appropriately classified all samples (Figure 2D). This model explains 17.7% of the variation in metabolite levels (R2X) and 76.9% of the variations between the different groups (R2Y). The average prediction capability (Q2) was 58%. The difference between R2Y and Q2 was less than 0.2 and the Q2 value was greater than 50%, revealing an excellent predictive capability (Robotti and Marengo, 2016). A 1,000-time permutation test was conducted to further validate the OPLS-DA model (Figure 2C). The empirical p-values for R2Y (p = 0.022) and Q2 (p < 0.001) indicate that the observed statistic (based on our data) is not part of the distribution formed by those from the permuted data. Thus, the dose-dependent clustering of the data is statistically significant.

**Figure 1.**
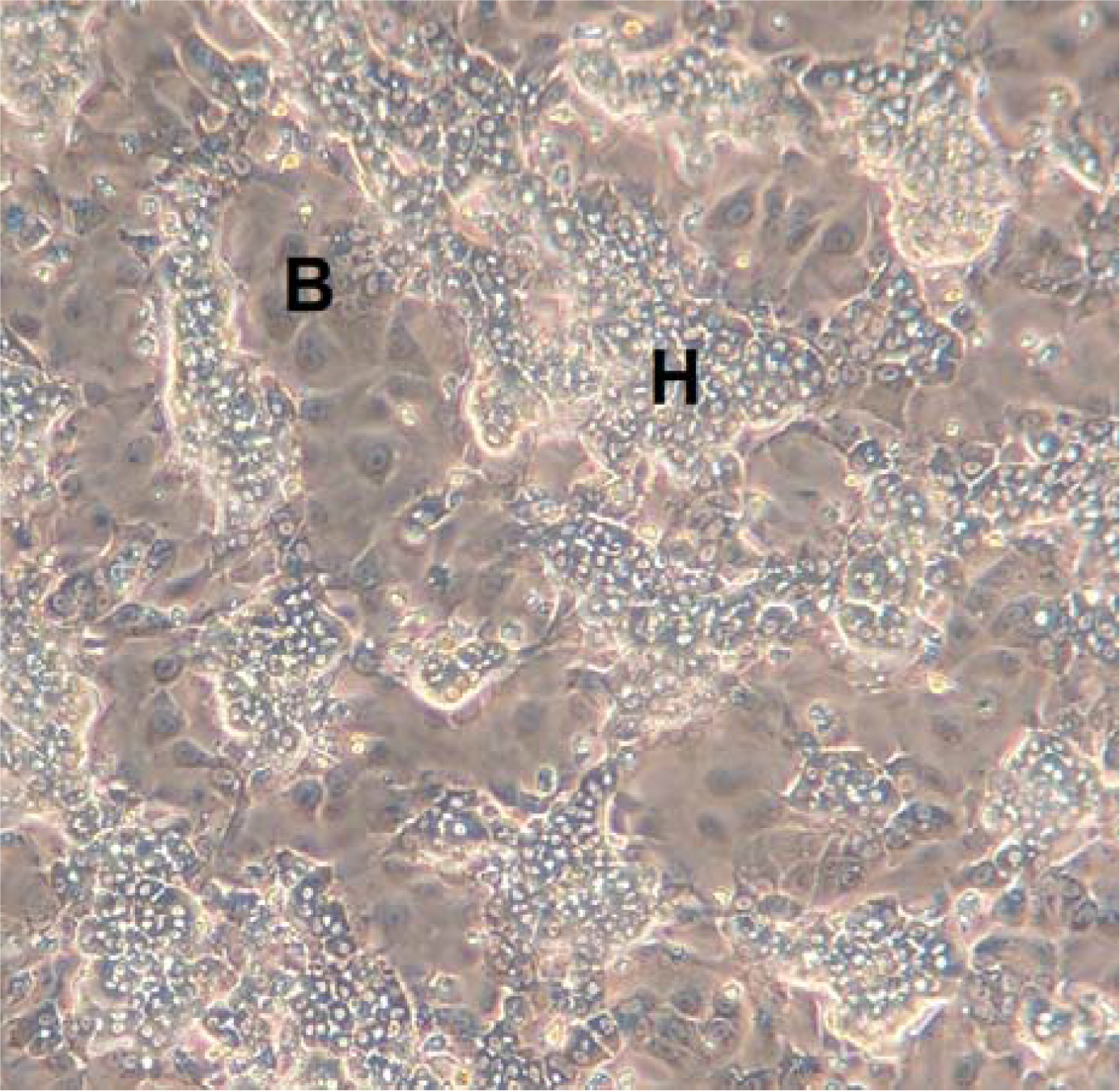
Morphology of HepaRG cells. After undergoing a complete programme of hepatocyte differentiation, HepaRG cells display the phenotype reflective of normal human liver cells including binuclear hepatocytes and forming bile canaliculus-like structures. A mixed population of 2 types of cells is visible, namely hepatocyte-like colonies (H) surrounded by clear epithelial cells corresponding to primitive biliary cells (B).

**Figure 2.**
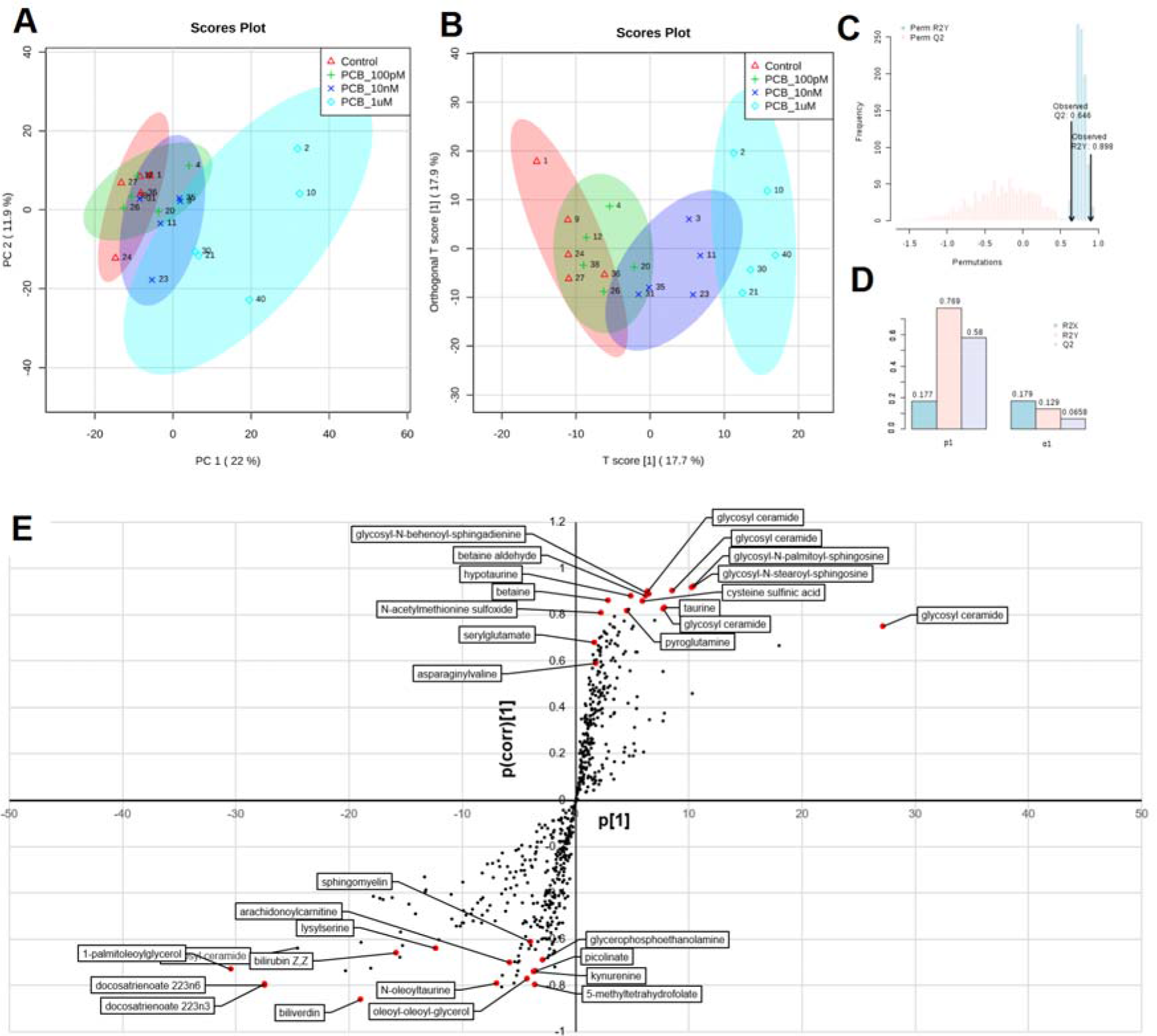
Multivariate analysis of HepaRG cell metabolome following treatment with PCB 126 shows alterations in lipid metabolism. **A**. Principal component analysis of metabolome profiles separate the group of samples treated with the PCB126 from the control group. As the dose of PCB 126 increases, the groups become more clustered. **B.** Orthogonal projection to latent structures discriminant analysis (OPLS-DA) properly classified all samples (R2X = 0.177, R2Y = 0.769, and Q2 = 0.58). The 95% confidence regions are displayed by shaded ellipses. **C.** A 1,000-time permutation test shows that the observed statistic is not part of the distribution formed by the statistic from the permuted data (R2Y p = 0.022; Q2 p < 0.001). **D**. Cross-validation parameters, R2 and Q2, representing the quality of the model **E.** The S-plot visualizes the variable influence in the OPLS-DA model. Significantly disturbed metabolites towards the separation in OPLS-DA models (red dots) were selected based on the significance threshold of q < 0.05 after analysis with one-way ANOVA test adjusted for multiple comparison with Fisher’s least significant difference. A total of 30 metabolites had their levels disturbed by the PCB 126 treatments.

An S-plot was then constructed to visualize the loading of the PCB 126 OPLS-DA model (Figure 2E), and determine which variables are the best discriminators between the different treatment groups. This plot combines the modelled covariance (p_1_) and the modelled correlation (p(corr)) from the OPLS-DA model in a scatter plot. Ceramides had the highest magnitude (p_1_) and reliability (p(corr)) scores. By contrast, the highest magnitude and reliability scores were attributable to polyunsaturated fatty acids (docosatrienoate 22:3n3 and 22:3n6). However, because OPLS-DA loading is difficult to interpret with more than 2 classes, we used the OPLS-DA method for score overview but for significantly disturbed metabolites (red dots, Figure 2E) selected based on the results of a one-way ANOVA test. A total of 30 metabolites had their levels disturbed by the PCB 126 treatments (Table 1). Most of the changes were dose dependent (Figure 3). The most pronounced metabolic differences were attributed to a change in lipid metabolism. A total of 4 glycosylceramides and 3 glycosyl-sphingosines had their levels increased (fold change (FC) up to 13.2 for glycosyl ceramide d181/231, d171/241), while polyunsaturated fatty acids such as docosatrienoate 22:3n3 (FC ranging from -1.3 to -5.4), and docosatrienoate 22:3n6 (FC from -1.6 to -6.5) and 1-palmitoleoylglycerol (FC from -1.8 to -370) had their levels decreased. These lipids are common constituents of cell membranes, and thus changes in their levels suggest that membrane integrity and maintenance is adversely affected by the PCB 126 treatment.

**Figure 3.**
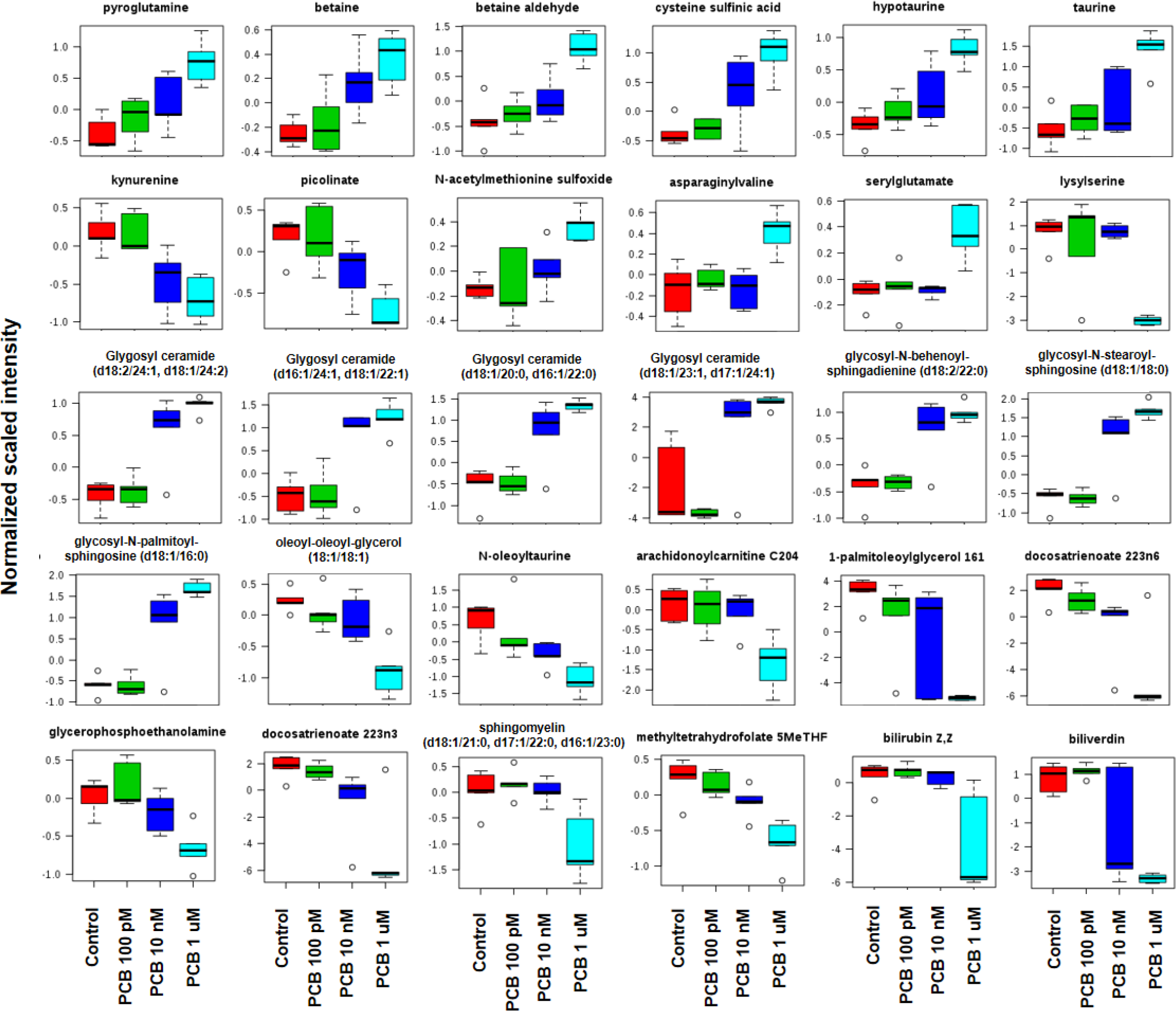
Box plot of metabolome changes associated with the exposure to PCB 126 to HepaRG cells shows a dose dependent effect. All the metabolites displayed have their levels significantly altered (q < 0.05) after analysis with one-way ANOVA test adjusted for multiple comparison with Fisher’s least significant difference. Most of the changes caused by the PCB treatment were dose dependent.

**Table 1.**
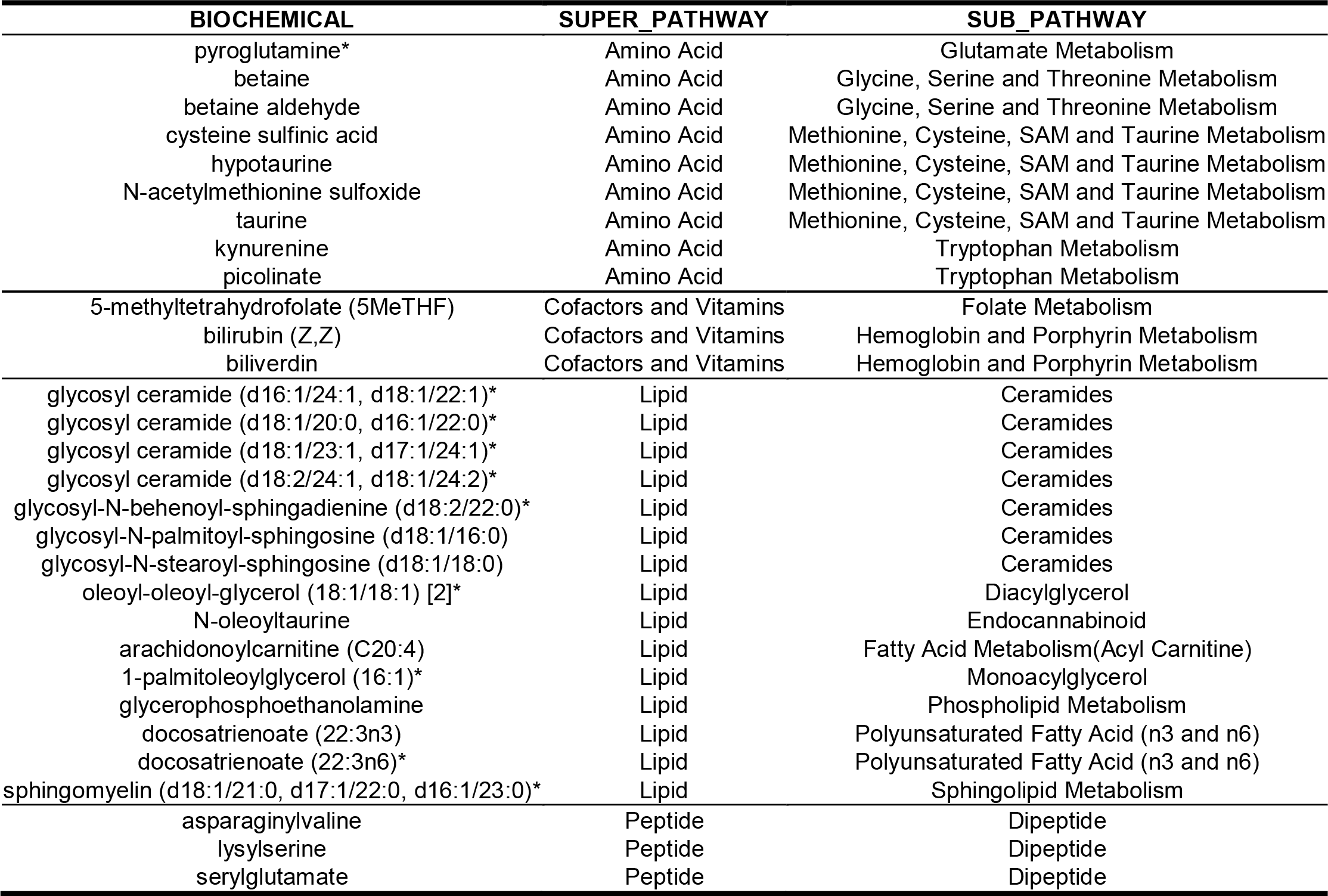
Metabolome disturbances provoked by exposure to PCB 126 in HepaRG cells. All the metabolites displayed have their levels significantly altered (q < 0.05) after analysis with one-way ANOVA test adjusted for multiple comparison with Fisher’s least significant difference.

Our results also show an increased activity within detoxification pathways. Multiple metabolites, including cysteine sulfinic acid, taurine, hypotaurine, N-oleoyltaurine, and N-acetylmethionine sulfoxide are significantly elevated in a dose-dependent manner. The predominant pathway elicited as a response to oxidative stress seems to be taurine metabolism as indicated by the network analysis performed using MetExplore (Figure 4A). Our network analysis reveals that bilirubin and biliverdin are further away from most metabolites in the fingerprint (Figure 4B). Thus, the network extraction focused on the first cluster which contains more metabolites (Figure 4A). The core part of the taurine and hypotaurine metabolism seems to be modulated (disregarding bilirubin and biliverdin) as a consequence of the oxidative stress induced by the PCB 126.

**Figure 4.**
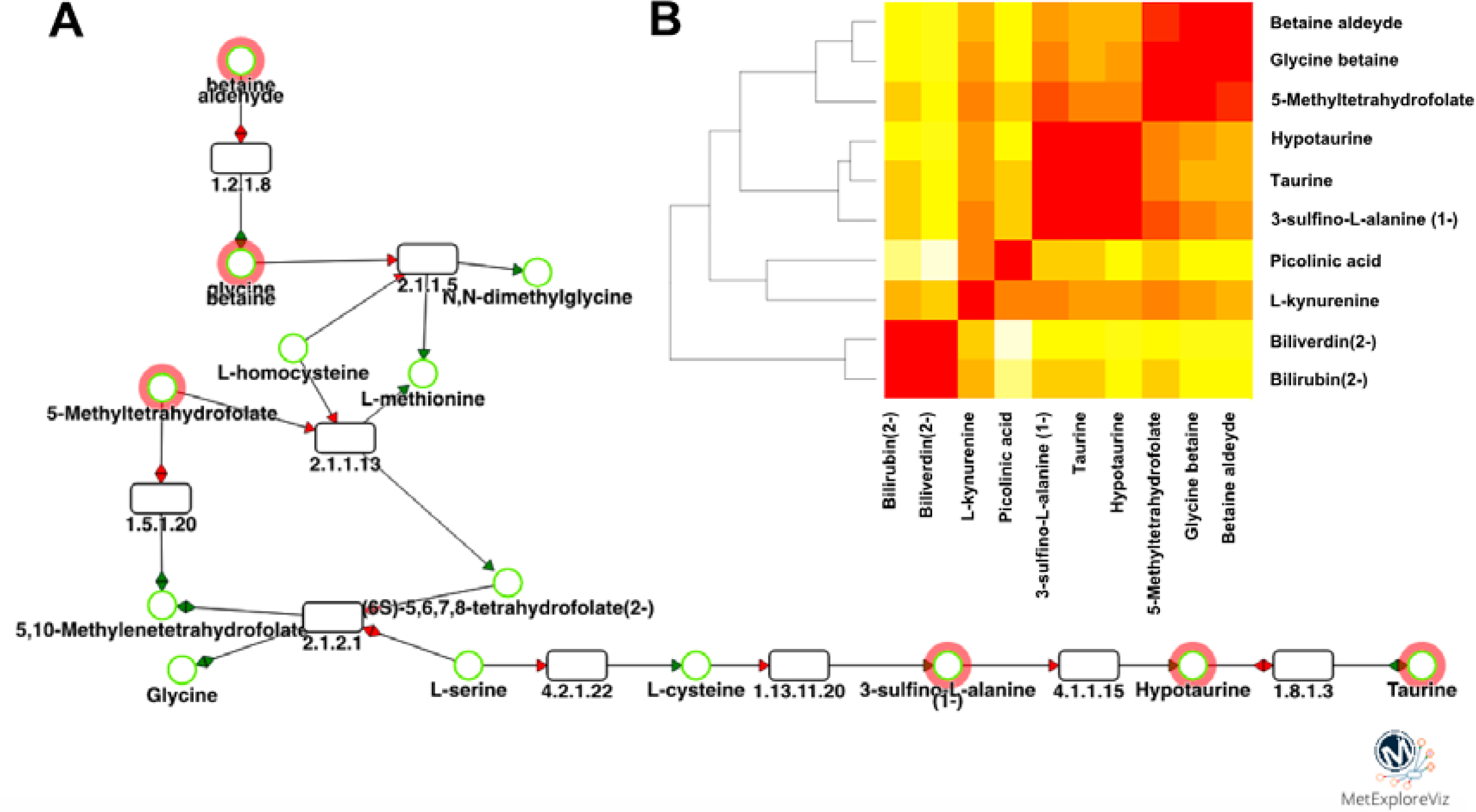
Network analysis of metabolome profile alterations demonstrates a role of taurine and hypotaurine in oxidative stress induced by PCB 126 in HepaRG cells. **A.** Sub-network. Circles are metabolites, rectangles are reactions. Reaction labels are EC numbers and metabolite names are from Recon2 model. Red circle metabolites are the ones from the fingerprint. **B.** Distance matrix between metabolites belonging to the network. Red corresponds to shorter distance (0) and white to longer distances (12 reactions between nodes).

Multiple gamma-glutamyl amino acids such as gamma-glutamylglutamine and gamma-glutamylcysteine are also elevated. This is possibly in response to oxidative stress being caused by the PCB 126 exposure. Several activators of aryl hydrocarbon receptor (AhR)-mediated transcription such as the bile pigments biliverdin and bilirubin, as well as tryptophan metabolites kynurenine and picolinate, had their levels decreased by the PCB 126 treatment. Other intermediates in tryptophan metabolism such as quinolinate, kynurenate and pyridoxate were also decreased but did not reach statistical significance.

Transcriptome profiles of HepaRG cells were then analysed using Illumina-based RNA sequencing. An unsupervised visualisation of the variance structure by a PCA showed comparable results to those of the metabolome profiles. The control group separates from the PCB 126 groups in a dose-dependent manner on the second component (Figure 5). The source of variation causing the clustering on the first component remains unknown as no batch effects was identified in the experimental procedure. As a result, the standard linear model-based comparison of transcript abundance was performed without adjusting for other covariates. A total of 264 transcripts had their levels altered by the PCB 126. We present a detailed analysis of *CYP1A1* expression as a representative of the changes we observed (Figure 6). *CYP1A1* is known to be highly expressed following exposure to dioxins and PCBs as its promoter contains 7 dioxin-responsive elements, which bind the aryl hydrocarbon receptor. The evaluation of expression levels among 12 distinct isoforms of *CYP1A1* shows that some transcripts are expressed at a much higher level than others (Figure 6A). Four of these 12 isoforms had their expression increased from the lowest concentration (q < 0.05). A scatter plot of the expression levels for the 24 most differentially expressed transcripts shows that alterations were dose-dependent and detectable at concentrations as low as 100 pM (Supplementary File 2).

**Figure 5.**
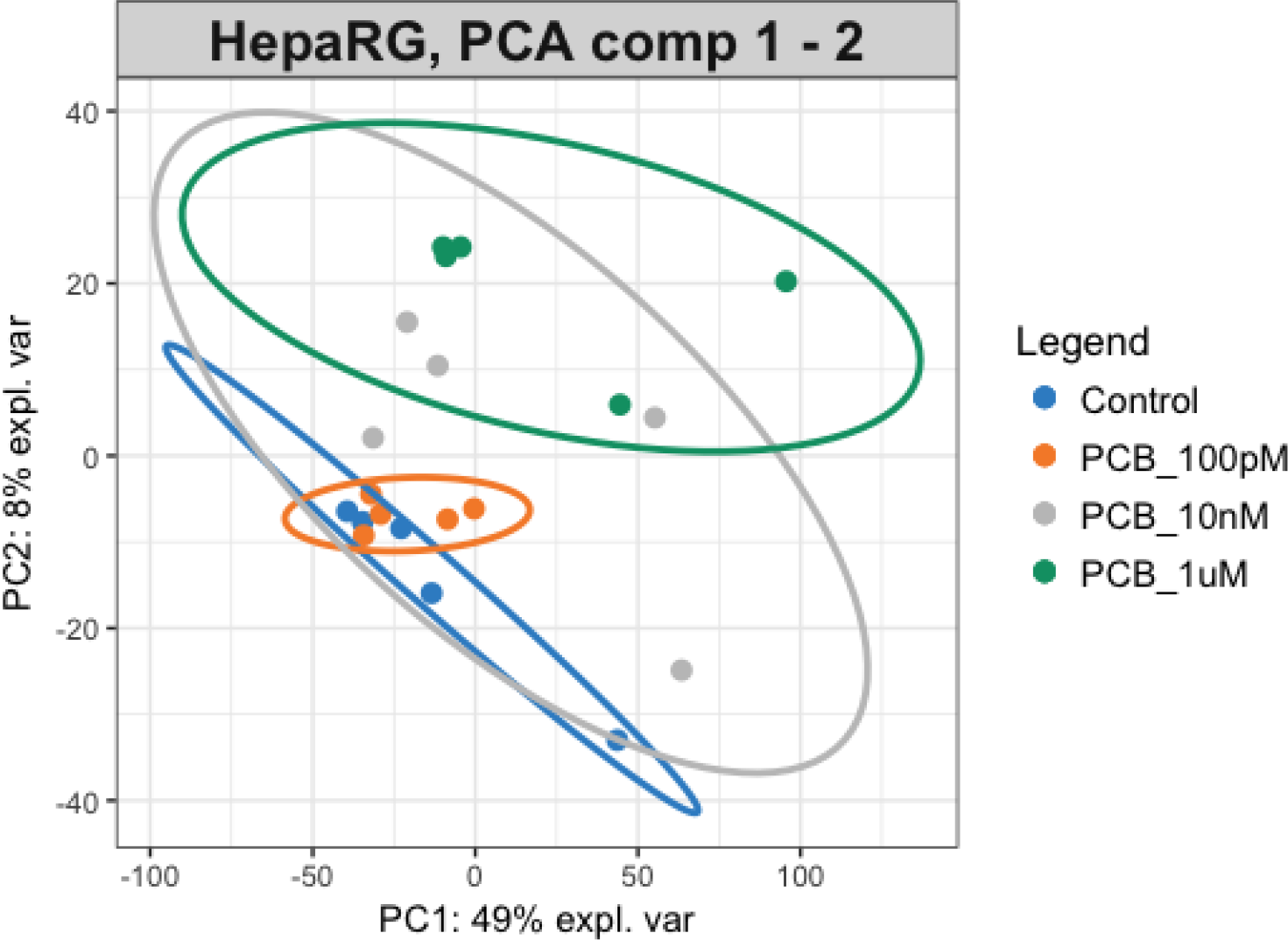
Principal component analysis of transcriptome profile alterations provoked by exposure of HepaRG cells to PCB 126. Transcript abundances were assessed using Stringtie. The PCA was performed using log2 transformed FPKM measurements of transcripts across samples assessed with Ballgown. The groups become more clustered as the dose of PCB 126 increases. The 95% confidence regions are displayed by ellipses.

**Figure 6.**
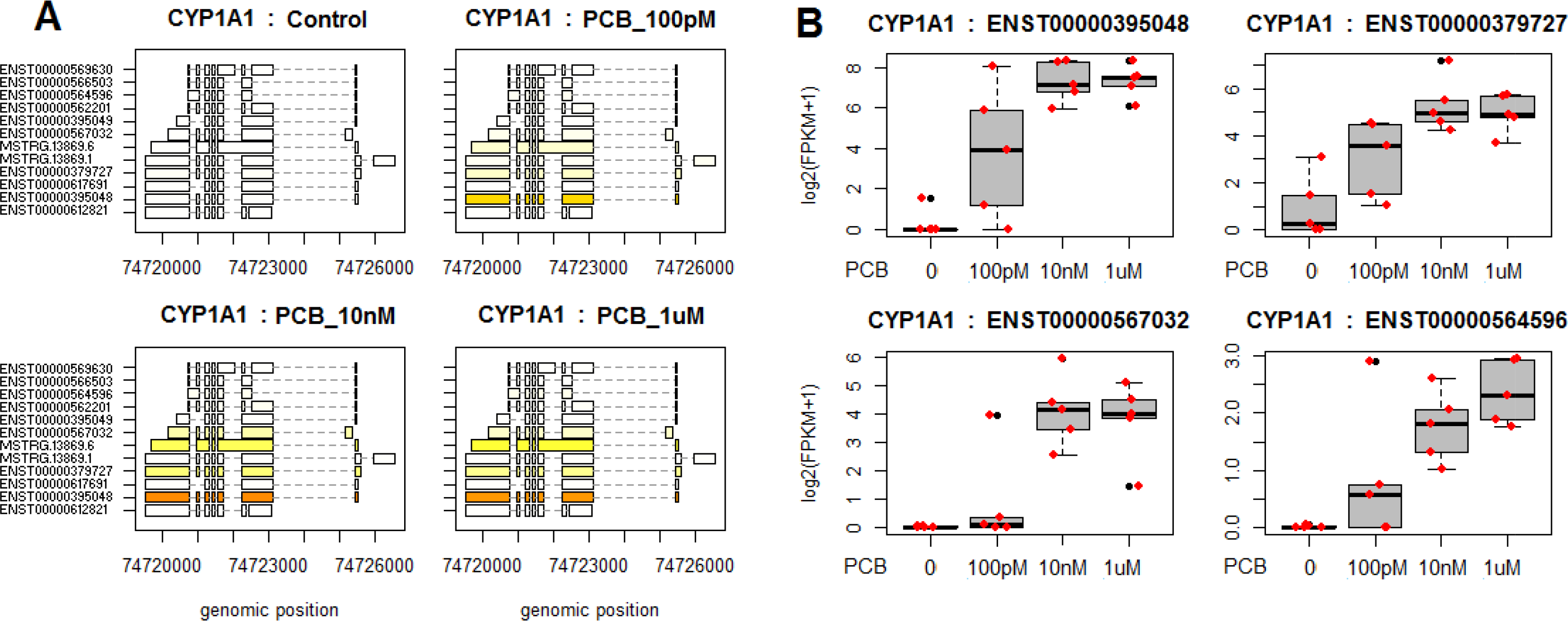
Differential *CYP1A1* expression analysis using RNA-seq in HepaRG cells exposed to three concentrations of PCB 126. **A**. Structure and expression levels of 12 distinct isoforms of *CYP1A1* across the different treatment groups. Differences in expression levels are displayed in varying shades of yellow. The ENST00000395048 isoform of *CYP1A1* is expressed at a much higher level than the others, as indicated by the dark orange colour. **B.** FPKM distributions of four *CYP1A1* transcripts displayed as box-and-whiskers plots. All four isoforms have their expression significantly altered (q < 0.05) by exposure to PCB 126 as measured by a standard linear model-based comparison in Ballgown.

We then performed a gene-annotation enrichment analysis in order to determine the biological processes affected by the exposure to PCB 126 (Figure 7, Supplementary File 3). Unsurprisingly, the most important network affected by PCB 126 were related to the metabolism of xenobiotics by P450 cytochromes (CYP1A1, CYP1A2, CYP1B1) and UDP glucuronosyltransferase (UGT1A1, UGT1A3, UGT2A3, UGT2B11), both being well known PCB metabolism pathways, but also aldehyde dehydrogenases (ALDH1A3, ALDH3A1), which catalyze oxidation of aromatic aldehydes produced by-products of mono-oxygenation reactions. These are hallmarks of an activation of xenobiotic metabolism by dioxin-like compounds (Grimm et al., 2015). More surprisingly, genes involved in the glycosaminoglycan metabolic process were the most enriched category (AP2A1, CEMIP, CHSY1, FUCA1, GALNT5, GPC6, LUM). This can reflect an increase in glucuronidation metabolism as UGT enzymes are central players in glycosylation processes. Perhaps one of the most surprising enriched pathways is the systemic lupus erythematosus (SLE) KEGG pathway containing histone genes (C5, H2AFV, HIST1H2AB, HIST1H2AE, HIST1H2AG, HIST1H2BO). The AGE-RAGE signalling pathway associated with diabetic complications was also enriched (AGTR1, CDKN1B, COL3A1, MMP2, PRKCA, SERPINE1). The same genes are found in the HIF-1 signalling pathway, which were also enriched. Altogether, these results point to a role of hypoxia in the toxic effects generated by the exposure to the PCB 126.

**Figure 7.**
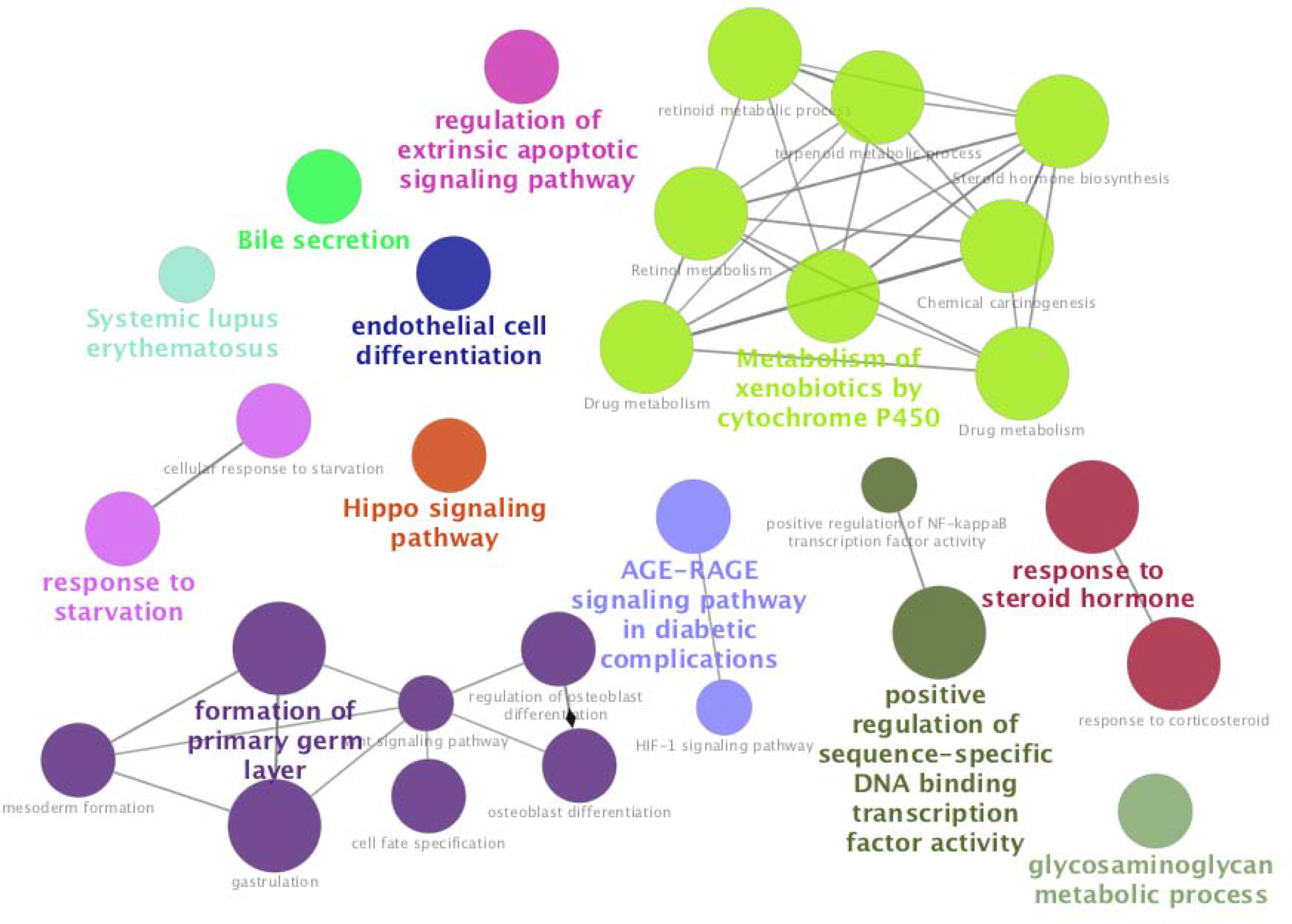
Pathway enrichment analysis in the transcriptome of HepaRG cells exposed to PCB 126 shows an activation of xenobiotic metabolism by dioxin-like compounds. Gene functions were studied using ClueGO and CluePedia plugins in Cytoscape (version 3.5.1). The analysis was conducted using a two-sided hypergeometric test for enrichment using a p-value threshold of 0.05 after its adjustment by the Benjamini-Hochberg procedure.

We then integrated the results from the transcriptome and the metabolome by performing an sPLS analysis, with two components being sufficient to model the data (Q2 of 0,1 and 0,17). The performance of the sPLS analysis was weak because of the relatively small number of samples. However, the different groups were well separated (Figure 8A), with the samples from the two highest concentrations of PCB 126 treatments being clustered together and separate from the control and the lowest dose samples. We then investigated the correlation between variables (Figure 8B and 8C). Sphingolipids and polyunsaturated fatty acids (PUFA) were negatively and positively correlated respectively with the expression of genes involved in triglyceride metabolism (such as APOA2, CPS1, HMGCS2, LPL). That is, the decrease in PUFA levels, as well as the increase in sphingolipid levels, is concomitant to a decrease in the activity of genes involved in lipid metabolism.

**Figure 8.**
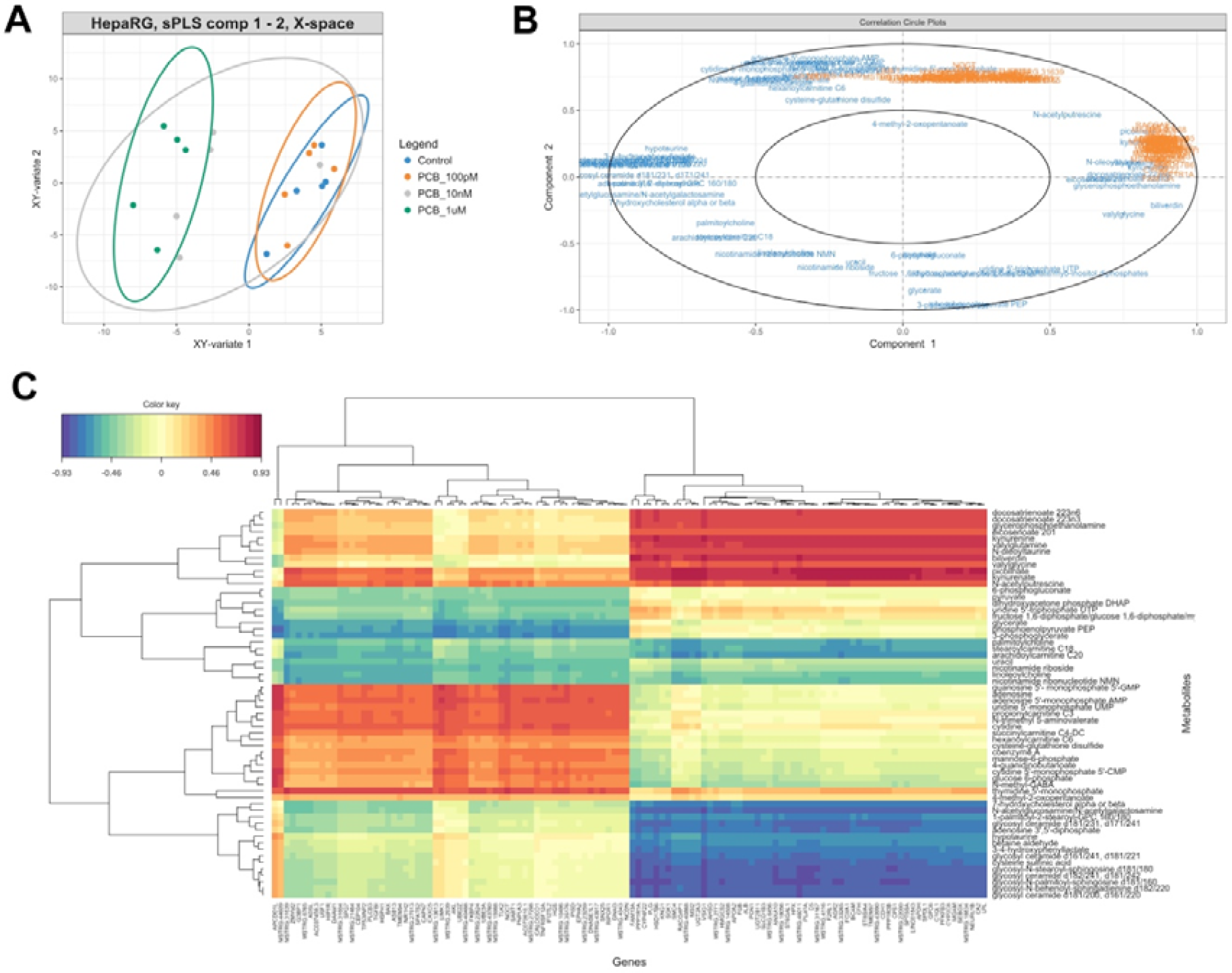
Sparse Partial Least Squares regression (sPLS) integration of the metabolome and transcriptome profiles of HepaRG cells exposed to PCB 126 shows that alteration in sphingolipid levels is concomitant to a decrease in the activity of genes involved in lipid metabolism. **A.** Individual plots displaying the covariance between the metabolome and the transcriptome datasets. **B.** The variables selected by the sPLS are projected on a correlation circle plot in order to display the clusters of correlated variables. In this plot, the angle defined by the coordinates of the variables on the axis defined by the components give an indication on the nature of the correlation. If the angle is sharp and the variables cluster together, the correlation is positive. If the angle is obtuse and the variables are not clustered together, the variables are negatively correlated. Perpendicular angles represent uncorrelated variables. **C.** A clustered image map visualises correlations between the metabolites and the genes by a colour gradient on a 2-dimensional colored image. The negatively correlated variables (blue) are represented along the positively correlated variables (red). Dendrograms are added to represent the clusters produced by the hierarchical clustering.

## Discussion

With the steadily increasing number of synthetic chemicals being released in our environment, the ability to use human cell lines to evaluate toxicity of pollutants and move away from more time-consuming and expensive risk assessment based on *in vivo* animal experiments, has become one of the challenges of the 21^st^ century (Hartung, 2009). We present here an in-depth investigation of the toxic effects of the PCB 126, a model chemical for the induction of TAFLD, on the transcriptome and metabolome of human liver HepaRG cells (Merrick et al., 2015; Mueller et al., 2014). We envisages that our investigation would be particularly useful for the establishment of high-throughput assays using the HepaRG cell line, which has been found to be a model system to predict the results of acute and repeated dose toxicity (Mueller et al., 2014) and to study human xenobiotic metabolism (Guillouzo et al., 2007). HepaRG is one of the best characterized cell lines. The most recent application of metabolomics analytical methods on the HepaRG cells reported coverage of 2200 metabolites (Cuykx et al., 2017). HepaRG cells are also metabolically competent and more stable than other hepatic cell lines, producing more reproducible data than primary human hepatocytes (Guillouzo et al., 2007). They are thus a reliable system to obtain information on networks of pathways adversely affected by pollutants that could lead to hepatic steatosis (Angrish et al., 2017).

Our study provides interesting insights into the development of sensitive biomarkers for TASH development. The most significant effect of PCB 126 exposure was a decrease in free long chain fatty acids (LCFAs) and PUFAs. Our results show that docosatrienoate 22:3n3 and 22:3n6 are the most sensitive biomarkers of lipid metabolism disturbances in HepaRG cells (Figure 1). This is in accord with the results obtained from a birth cohort from a fishing community that linked a high seafood intake, and thus high PCB exposure, to alterations in lipid profiles (Grandjean and Weihe, 2003). Other experimental studies suggest that PCBs may affect the utilization of PUFAs by inhibition of Δ5- and Δ6-desaturation (Matsusue et al., 1999). Such enzyme inhibition in fishing communities lead to deficient formation of arachidonic acid from its precursor, linoleic acid, attenuating the beneficial effects of essential lipids contained in seafood (Grandjean and Weihe, 2003).

The analysis of gene expression profiles revealed alterations in pathways reflecting PCB-induced metabolic diseases such as the activation of xenobiotic metabolism by dioxinlike compounds. This can guide the development of gene expression biomarkers, which have been advocated by the US Environmental Protection Agency (EPA) as promising tools to reduce the cost and time of toxicity testing (EPA., 2015), such as for the detection of potential EDCs acting as estrogen receptor agonists (Mesnage et al., 2017). Other projects have implemented online tools to provide an easy access to gene expression signatures allowing the detection of novel associations between genes, diseases and drugs (Wang et al., 2016). We provide here putative biomarkers of metabolic disturbances induced by PCB 126, which are known to correlate with the mechanisms by which this class of pollutants induce TASH (Al-Eryani et al., 2015). The expected hallmarks of an activation of xenobiotic metabolism by dioxin-like compounds were detected in the molecular profiles of HepaRG cells exposed to the PCB 126 (Figure 7). A limit of this study is that the changes induced by the PCB 126 were measured at a late time point and may not reflect the first wave of transcriptional changes induced by the PCB 126.

Other pathways altered in our study could correlate with other less known health effects of PCB compounds. For instance, SLE KEGG pathway was one of the most altered. This in all likelihood is not coincidental as the exposure to PCBs has been linked to an increased incidence of SLE (Kamen, 2014). Histone gene expression was the most disturbed of the SLE pathway in our study (Supplementary Material 3). This correlates with the fact that patients with SLE often carry anti-histone antibodies (Sun et al., 2008). A link with SLE is further corroborated by laboratory experiments showing that docosahexaenoic acid consumption attenuates the autoimmune response in a mouse model of SLE (Bates et al., 2016). It may be argued that the detection of this gene expression signature in an hepatic cell line has no direct relevance to SLE. However, comparable responses have been found in other cell types equipped with AhR signalling (Vorrink et al., 2014). Another category of genes whose levels we found altered were those involved in the glycosaminoglycan metabolic process. This correlates with findings obtained from the analysis of the gene ontology enrichment in peripheral blood mononuclear cells of children chronically exposed to PCBs (Dutta et al., 2012).

Our study also provides mechanistic insights into perturbations provoked by very low environmental concentrations of PCB 126. Although the exposure to persistent organic pollutants such as PCBs is decreasing overtime as they are progressively banned, they are still found in human tissue at levels capable of causing pathologies. PCB 126 was found at concentrations of 209 ng/kg and 80.4 ng/kg in the breastmilk of Inuit women and Caucasian women, respectively, in a survey performed in 1989-1990 (Dewailly et al., 1994). The analysis of the National Health and Nutrition Examination Survey (NHANES) data (1999-2002) reveals associations between PCB serum levels and adverse cognitive effects in older US adults (Przybyla et al., 2017). In more recent biomonitoring surveys, PCB 126 was found at a mean concentration of 6.5 µg/L in umbilical cord serum of populations living near waste dump sites in Italy (Grumetto et al., 2015).

The transcriptome and metabolome signatures that we have identified in response to PCB 126 in HepaRG cells are likely to be similar to what occurs in cases of NAFLD caused by other factors. It is known from metabolome profiles that hepatic PUFAs are lower in NAFLD patients (Arendt et al., 2015). This suggests that the exposure to environmental pollutants could also be an aggravating factor in the development of NAFLD, amplifying the effects of over-nutrition or a sedentary lifestyle. It has been proposed that PCB exposure can act as a second hit in NAFLD, driving progression of steatosis to NASH as has been found in laboratory animals exposed to PCB and a high fat diet (Wahlang et al., 2014).

## Conclusion

Although is it recognized that occupational exposures to chemical pollutants can lead to NAFLD, there is limited information available regarding the mechanism by which typical environmental levels of exposure can contribute to the onset of this disease. Importantly, the alterations in metabolome and transcriptome profiles in response to PCB 126 were observed even at the lowest concentration (100 pM) tested corresponding to what may be found in humans. Overall, our integrated multi-omics analysis provides mechanistic insight into how this class of chemical pollutant can cause NAFLD as demonstrated in other studies (Al-Eryani et al., 2015). In addition, we confirm the HepaRG cell line as a reliable model for molecular profiling investigations of TAFLD. Metabolome profiling reveals docosatrienoate levels as the most reliable marker of the exposure to PCB 126. Hallmarks of activation of the AhR receptor by dioxin-like compounds were detected in the transcriptome profiles. Our study provides the foundation for the development of molecular signatures of fatty liver diseases to rapidly assess chronic toxic effects in a reliable hepatoxicity model system, in order to lessen the burden of chemical risk assessment based on *in vivo* animal experiments.

## Acknowledgements

This work was funded by the Sustainable Food Alliance (USA) whose support is gratefully acknowledged.

## Authors’ contributions

RM participated in the cell culture experiment, performed the metabolome and transcriptome data analysis, and drafted the manuscript. MB performed the cell culture experiment and the RNA extraction. SB performed the library preparation. CF, NP and FJ performed the network analysis of the metabolome data. EW, TX and CAM performed the RNA-seq and assisted for the transcriptome data analysis. MNA coordinated the investigation and drafted the manuscript.

## Supplementary Data

**Supplementary File 1. Summary of RNA-seq alignment files.**

**Supplementary File 2. Box plot of transcriptome changes associated with the exposure to PCB 126 in HepaRG cells.** All the transcript displayed have their levels significantly altered (q < 0.05). Most of the changes caused by the PCB treatment were dose dependent.

**Supplementary File 3.** Gene ontology analysis performed with ClueGO. Functional implications of the alteration in gene expression profiles were analysed using ClueGO (Bindea et al., 2009) and CluePedia plugins in Cytoscape (version 3.5.1). The GO biological process database (23.02.2017) and the KEGG annotation database (01.03.2017) were used. The analysis was conducted using a two-sided hypergeometric test for enrichment using a p-value threshold of 0.05 after its adjustment by the Benjamini-Hochberg procedure. GO term fusion was employed to integrate GO categories, minimize the output, and create a functionally organized GO category network. Our network displayed GO terms found in the levels 5-10 of the GO hierarchy, in order to avoid meaningless high-level hierarchy terms.

